# Sphingolipid metabolism drives mitochondria remodeling during aging and oxidative stress

**DOI:** 10.1101/2025.02.26.640157

**Authors:** Adam C. Ebert, Nathaniel L. Hepowit, Thyandra A. Martinez, Henrik Vollmer, Hayley L. Singkhek, Kyrie D. Frazier, Sophia A. Kantejeva, Maulik R. Patel, Jason A. MacGurn

## Abstract

One of the hallmarks of aging is a decline in the function of mitochondria, which is often accompanied by altered morphology and dynamics. In some cases, these changes may reflect macromolecular damage to mitochondria that occurs with aging and stress, while in other cases they may be part of a programmed, adaptive response. In this study, we report that mitochondria undergo dramatic morphological changes in chronologically aged yeast cells. These changes are characterized by a large, rounded morphology, decreased co-localization of outer membrane and matrix markers, and decreased mitochondrial membrane potential. Notably, these transitions are prevented by pharmacological or genetic interventions that perturb sphingolipid biosynthesis, indicating that sphingolipids are required for these mitochondrial transitions in aging cells. Consistent with these findings, we observe that overexpression of inositol phospholipid phospholipase (Isc1) prevents these alterations to mitochondria morphology in aging cells. We also report that mitochondria exhibit similar sphingolipid-dependent morphological transitions following acute exposure to oxidative stress. These findings suggest that sphingolipid metabolism contributes to mitochondrial remodeling in aging cells and during oxidative stress, perhaps as a result of damaged sphingolipids that localize to mitochondrial membranes. These findings underscore the complex relationship between mitochondria function and sphingolipid metabolism, particularly in the context of aging and stress.

## Introduction

Declining function of mitochondria is associated with many models of aging and disease and correlates with striking morphological changes. In healthy cells, mitochondria are long, tubular structures, while in aging or stressed cells they become fragmented or swollen (Chaudhari and Kipreos, 2017; Houtkooper et al., 2013; Lima et al., 2023). For example, aged cells in a yeast replicative aging model have fragmented mitochondria with decreased inner membrane potential (Hughes and Gottschling, 2012). Mitochondrial swelling has been reported in human cardiomyocytes following ischemic injury, in motor neurons of aged rats and in mitochondria of aging worms (García et al., 2013; Li et al., 2023; Rosa et al., 2023). These morphological changes are often associated with altered mitochondrial dynamics, and interventions blocking fission or promoting fusion can preserve mitochondria in aging cells and increase lifespan (Chaudhari and Kipreos, 2017; Scheckhuber et al., 2007). However, little is known about the factors that drive these transitions in aging mitochondria.

Sphingolipids (SLs) are synthesized in the endoplasmic reticulum (ER) and Golgi and contribute to membrane structures throughout the cell. The first step of SL biosynthesis is highly conserved in eukaryotic cells and involves condensation of serine and palmitoyl CoA by the serine palmitoyltransferase (SPT) enzyme in the ER. The product is further modified and acylated to produce ceramide, which can then be transported to the Golgi and modified to more complex SLs. While biosynthesis of ceramides in the ER is conserved across eukaryotes, modifications in the Golgi are more divergent. For example, the yeast *S. cerevisiae* attaches inositol head groups to ceramides which are further modified to more complex SLs, while in animal cells complex SLs are derived from attachment of either glucose or phosphocholine head groups to produce gangliosides or sphingomyelins, respectively. These complex SLs contribute to the structure of cellular membranes and also have important signaling and regulatory functions in the cell.

Accumulating evidence suggests that sphingolipids (SLs) are detrimental to mitochondrial health, which may explain why SL depletion promotes longevity and healthspan (Cutler et al., 2014; Hepowit et al., 2021; Huang et al., 2014; Li and Kim, 2021; Singh and Li, 2018). SL depletion improved mitochondrial function in aged mice and in human primary muscle cells from aged donors (Lima et al., 2023). In mice, a ceramide synthase (CerS6) is required for obesity-induced mitochondrial fragmentation and remodeling, and loss of this enzyme protects mice from diet-induced obesity (Hammerschmidt et al., 2019). Ceramides were found to accumulate in mitochondria of aged *parkin* null mice, a model for Parkinson’s Disease (PD) (Gaudioso et al., 2019) as well as *pink1* mutant flies and patient-derived fibroblasts (Vos et al., 2021). In *pink1* mutant flies, SL depletion rescued mitochondria morphology, ETC function, and flying ability (Vos et al., 2021). In a *C. elegans* model of Aβ proteotoxicity, SL depletion rescued mitochondrial defects (Lima et al., 2023). All of these results suggest that SL accumulation can be a contributing factor to mitochondria dysfunction in states of disease, but it remains unclear why and how sphingolipids are a liability for mitochondrial function.

Here, we report that mitochondria undergo dramatic morphological transitions in a yeast cell model of chronological aging. These transitions are sphingolipid-dependent, and interventions that deplete sphingolipids preserve mitochondria and increase longevity. Notably, we observe a similar relationship between sphingolipid metabolism and mitochondrial morphology during the cellular response to acute oxidative stress. These findings underscore the close relationship between sphingolipid metabolism and mitochondrial health and may help explain why sphingolipid depletion is associated with increased longevity in various aging models.

## Results and Discussion

### Mitochondria undergo morphological changes during chronological aging

To characterize how mitochondria morphology changes in aging cells, we analyzed yeast expressing Tom70-mCherry (to label the outer mitochondrial membrane (OMM) and Mdh1-GFP (to label the mitochondrial matrix (MM)) at different time points during a chronological aging time course. We observed that mitochondria transition from filamentous to fragmented after exiting mitosis (**Fig 1A** and **FIG S1A**, 24 hour time point) and as cells age the mitochondria transition to rounded structures (**FIG 1A** and **FIG S1A,** 48 and 72 hour time points). These age-associated changes in mitochondria morphology are characterized by increased width (**FIG 1B**), decreased co-localization between Tom70-mCherry and Mdh1-GFP (**FIG 1C**) and decreased fluorescence intensity signal from Mdh1-GFP (**FIG 1D**). These features of chronological aging were observed using different strain backgrounds (**FIG 1A-D** and **FIG S1A-C**) and using different markers for the OM or MM (**FIG S1D**). Notably, we observed these transitions in different defined, minimal media (e.g., SCD and Longo’s) but in rich media (YPD) mitochondria exhibited fragmentation but did not exhibit rounding by 48 hours of aging (**FIG S1E-F**).

**Figure 1.**
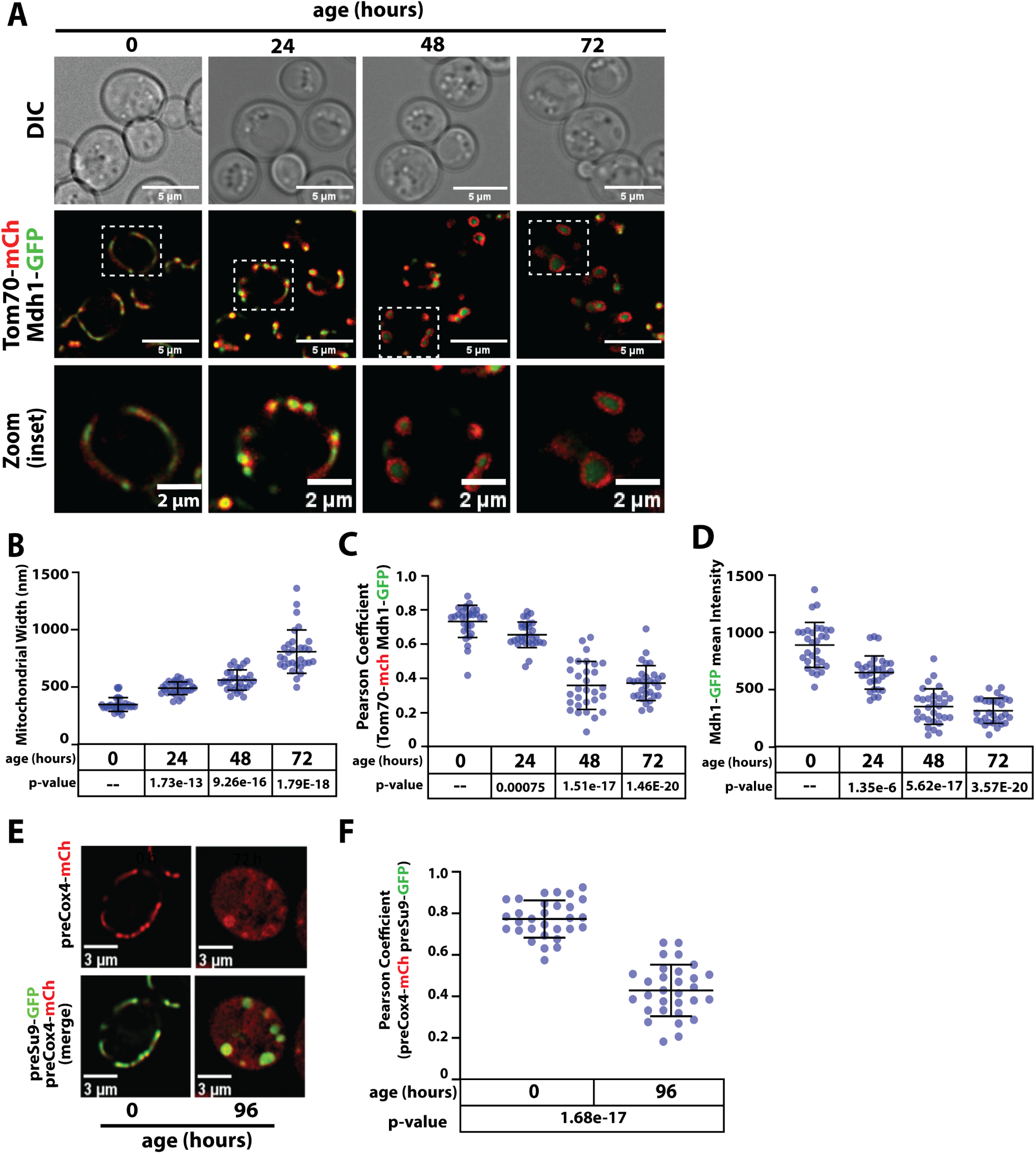
Yeast chronological aging is associated with mitochondrial remodeling and functional decline. (A) Yeast cells (strain background: SEY6210) expressing Tom70-mCherry (red) and Mdh1-GFP (green) were grown to mid-log phase (OD_600_ = 0.5) to initiate chronological aging time course (t=0). Fluorescence microscopy images were collected every 24 hours at the indicated time points. (B) Mitochondria width was measured (n ≥ 30 cells; ±SD (error bars)) at each time point. The indicated p-value was computed using a paired T-TEST comparing the indicated time point to t=0. (C) Pearson correlation coefficient was computed for individual cells (n ≥ 30 cells; ±SD (error bars)) using Tom70-mCherry (red) and Mdh1-GFP (green) as markers of the outer mitochondrial membrane (OMM) and mitochondrial matrix (MM), respectively. The indicated p-value was computed using a paired T-TEST comparing the indicated time point to t=0. (D) The total cellular intensity of Mdh1-GFP was computed for individual cells (n ≥ 30 cells; ±SD (error bars)) at each time point. The indicated p-value was computed using a paired T-TEST comparing the indicated time point to t=0. (E) Mitochondrial membrane potential (MMP) was assessed using the MitoLoc system. This assay relies on use of two mitochondrial import reporters: preSu9-GFP (green), which localizes to mitochondria independent of MMP, and preCox4-mCherry (red) which localizes to mitochondria in a MMP-dependent manner. Thus, accumulation of mCherry signal outside of mitochondria indicates loss of MMP. (F) Pearson correlation coefficient was computed for individual cells (n ≥ 30 cells; ±SD (error bars)) using preCox4-mCherry (red) and preSu9-GFP (green) of the MitoLoc system. The indicated p-value was computed using a paired T-TEST comparing the indicated time point to t=0.

To assess mitochondrial membrane potential (MMP), we used the MitoLoc reporter system (Vowinckel et al., 2015). MitoLoc relies on two fluorescent reporters: preSu9-GFP, which is imported to the MM independent of MMP, and preCox4-mCherry, which is imported to the MM in an MMP-dependent manner (Vowinckel et al., 2015). We observed less Cox4-mCherry localization in mitochondria of aged cells, consistent with decreased MMP (**FIG 1E-F**).

Decreased MMP was also reported for mitochondria in the yeast replicative aging model (Hughes and Gottschling, 2012) although the morphological changes that occur during replicative aging appear distinct. Since mitochondria decline in replicative aging can be suppressed by enhancing vacuolar function (Hughes and Gottschling, 2012) we tested if mitochondria morphology during chronological aging can likewise be suppressed by enhancing vacuolar function. We found that overexpression of factors that preserve mitochondria in a replicative aging model (*AVT1*, *VMA1*, and *VPH2* (Hughes and Gottschling, 2012)) did not prevent changes to mitochondrial morphology during chronological aging (**FIG S1G-H**).

Furthermore, while Tom70 was reported to localize in vacuoles in a replicative aging model (Hughes et al., 2016) we did not detect significant vacuolar localization of Tom70 in a chronological aging model (**FIG S1I-J**). Taken together, our results indicate that mitochondria undergo morphological changes and functional decline during chronological aging by a mechanism that is distinct from replicative aging.

### Myriocin preserves mitochondria in chronologically aged cells

To better understand the mechanisms underlying these morphological changes, we tested if interventions known to enhance longevity preserve mitochondria morphology during chronological aging. Several compounds associated with increased longevity in yeast (e.g., ibuprofen and mycophenolic acid (Sarnoski et al., 2017)) and other organisms (e.g., taurine (Singh et al., 2023)) did not affect mitochondria morphology during chronological aging.

However, we found that both rapamycin (rap; an inhibitor of TORC1) and myriocin (myr; which inhibits the first step of sphingolipid (SL) biosynthesis) prevented mitochondrial morphology changes during chronological aging, although neither compound prevented mitochondrial fragmentation observed at earlier time points (**FIG 2A-B**). We also found that lowering glucose concentration in the media, which is associated with increased longevity by caloric restriction (Guo et al., 2022; Zou et al., 2020), prevented mitochondria morphology changes associated with chronological aging (**FIG S2A-B**). Together, these results indicate that a subset of longevity-associated interventions, including caloric restriction, TORC1 inhibition, and sphingolipid depletion, can preserve mitochondria in chronologically aged cells.

**Figure 2.**
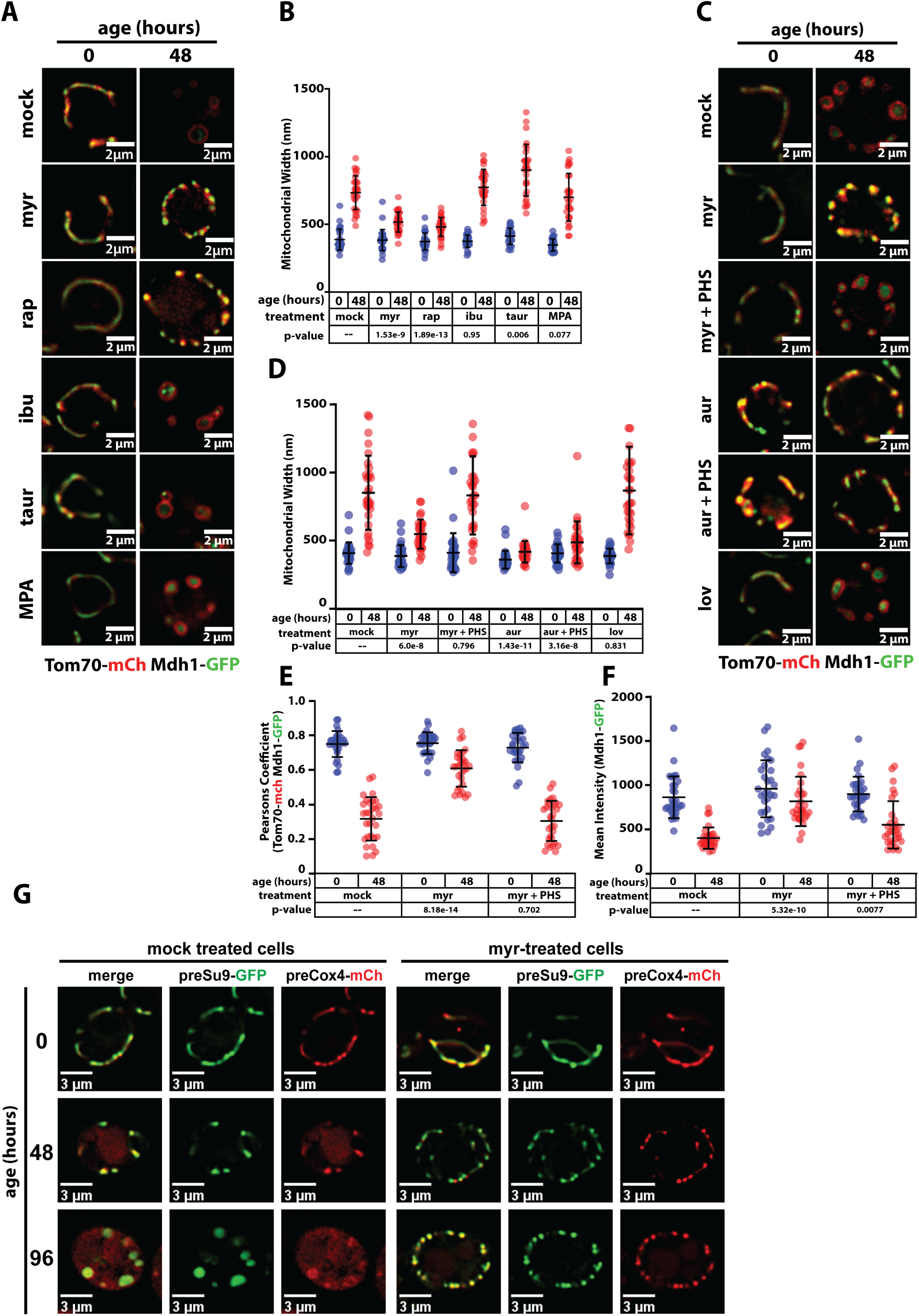
Myriocin preserves mitochondrial morphology and function in aged yeast cells. (A) Yeast expressing Tom70-mCherry (red) and Mdh1-GFP (green) were grown to mid-log phase (OD_600_ = 0.5) to initiate chronological aging time course (t=0). Fluorescence microscopy images were collected again after 48 hours of aging. Cells were left untreated (mock) or treated with the following compounds associated with increased life span: myriocin (myr), rapamycin (rap), ibuprophen (ibu), taurine (taur) and mycophenolic acid (MPA). (B) Mitochondria width was measured (n ≥ 30 cells; ±SD (error bars)) for each condition shown in (A). The indicated p-value was computed using a paired T-TEST comparing the indicated treatment (t=48) to mock treated cells (t=48). (C) Yeast expressing Tom70-mCherry (red) and Mdh1-GFP (green) were grown to mid-log phase (OD_600_ = 0.5) to initiate chronological aging time course (t=0). Fluorescence microscopy images were collected again after 48 hours of aging. Cells were left untreated (mock) or treated with the following compounds that perturb lipid metabolism: myriocin (myr), myriocin + phytosphingosine (myr + PHS), aureobasidin A (aur), aureobasidin A + phytosphingosine (aur + PHS), and lovastatin (lov). (D) Mitochondria width was measured (n ≥ 30 cells; ±SD (error bars)) at each time point. The indicated p-value was computed using a paired T-TEST comparing the indicated treatment (t=48) to mock treated cells (t=48). (E) Pearson correlation coefficient was computed for individual cells (n ≥ 30 cells; ±SD (error bars)) using Tom70-mCherry (red) and Mdh1-GFP (green) as markers of the outer mitochondrial membrane (OMM) and mitochondrial matrix (MM), respectively. The indicated p-value was computed using a paired T-TEST comparing the indicated treatment (t=48) to mock treated cells (t=48). (F) The total cellular intensity of Mdh1-GFP was computed for individual cells (n ≥ 30 cells; ±SD (error bars)) at each time point. The indicated p-value was computed using a paired T-TEST comparing the indicated treatment (t=48) to mock treated cells (t=48). (G) Mitochondrial membrane potential (MMP) was assessed using the MitoLoc system. This assay relies on use of two mitochondrial import reporters: preSu9-GFP (green), which localizes to mitochondria independent of MMP, and preCox4-mCherry (red) which localizes to mitochondria in a MMP-dependent manner. Thus, accumulation of mCherry signal outside of mitochondria indicates loss of MMP.

We were interested to better understand the relationship between sphingolipid metabolism and mitochondrial morphology in aging cells. We found that both myriocin (Myr) and aureobasidin A (AurA) prevented the transition to larger, rounded mitochondria in chronologically aged cells (**FIG 2C-F**). The effect of Myr is suppressed by adding phytosphingosine (PHS) to the media which bypasses the pharmacological blockade, confirming SL-dependence of the age-associated changes in mitochondrial morphology (**FIG 2C-F**). In contrast, PHS does not suppress the effect of AurA (**FIG 2C-D**) indicating a requirement for synthesis of complex SL species downstream of the enzyme Aur1, which adds an inositol head group to ceramide (see **FIG 3A**). Preservation of mitochondria morphology by rapamycin is not suppressed by PHS supplementation (**FIG S2C-D**). Notably, lovastatin, which inhibits synthesis of sterols, did not prevent mitochondria morphology changes associated with chronological aging (**FIG 2C-D**).

Myriocin treatment also preserved MMP in chronologically aging cells (**FIG 2G**) indicating that both morphology changes and functional decline of mitochondria are linked to sphingolipid levels.

### Complex SL biosynthesis is required for age-associated mitochondrial transitions

To further explore the relationship between SL biosynthesis (**FIG 3A**) and mitochondrial health, we analyzed mitochondrial aging in cells lacking non-essential SL biosynthesis enzymes (**FIG 3B**). Various strains deleted for SL biosynthesis enzymes, including Sur2 (which converts dihydrosphingosine (DHS) to phytosphingosine (PHS)), Lac1 (a subunit of the ceramide synthase complex), and Csg2 (part of the MIPC synthase complex)) exhibited mitochondrial fragmentation but not the rounded mitochondrial morphology observed in aging wildtype cells (**FIG 3B-C**). Notably, supplementation of growth media with PHS restored wildtype mitochondrial morphology in aging *Δsur2* cells, but not in *Δlac1* or *Δcsg2* cells (**FIG 3B-C**). These data indicate that complex SLs downstream of IPC are required for the mitochondrial morphology observed in chronologically aging cells.

**Figure 3.**
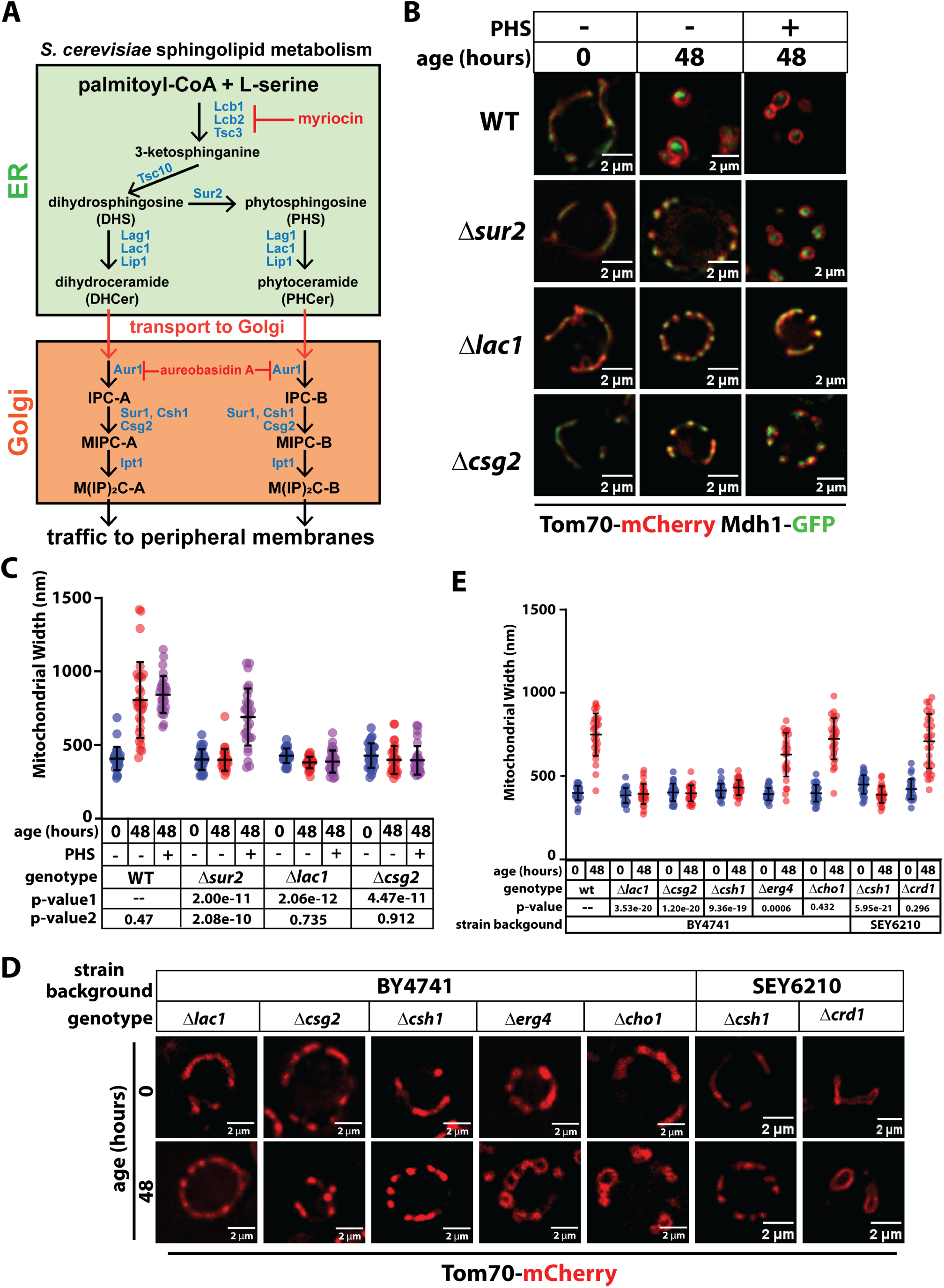
Sphingolipid biosynthesis is required for age-associated changes to mitochondria morphology. (A) Diagram of the sphingolipid (SL) biosynthesis pathway in yeast. Key enzymes in this pathway are colored in blue, while inhibitors used in this study are colored in red. (B) Yeast expressing Tom70-mCherry (red) and Mdh1-GFP (green) were grown to mid-log phase (OD_600_ = 0.5) to initiate chronological aging time course (t=0). Fluorescence microscopy images were collected again after 48 hours of aging. Wildtype (WT) cells were analyzed as well as the indicated knockout mutants. PHS was added to supplement the media where indicated. (C) Mitochondria width was measured (n ≥ 30 cells; ±SD (error bars)) for each condition shown in (B). The p-value 1 was computed using a paired T-TEST comparing the indicated mutant cells (t=48) to wildtype cells (t=48). The p-value 2 was computed using a paired T-TEST comparing the indicated mutant cells (t=48) to the same mutant cells grown in media supplemented with PHS (t=48). (D) Yeast expressing Tom70-mCherry were grown to mid-log phase (OD_600_ = 0.5) to initiate chronological aging time course (t=0). Fluorescence microscopy images were collected again after 48 hours of aging. Wildtype (WT) cells were analyzed as well as the indicated knockout mutants. Two different strain backgrounds were used in this analysis: BY4741 and SEY6210. (E) Mitochondria width was measured (n ≥ 30 cells; ±SD (error bars)) for each condition shown in (D). The p-value was computed using a paired T-TEST comparing the indicated mutant cells (t=48) to wildtype cells (t=48).

We next expanded our analysis to include different lipid biosynthesis mutants available in yeast knockout collection (BY4741 strain background). Consistent with previous experiments, mutants disrupted for SL biosynthesis (*Δlac1*, *Δcsg2*, *Δcsh1*) did not exhibit rounded mitochondria in aging cells, again revealing that complex SLs are required for aging mitochondria morphology (**FIG 3D-E**). In contrast, morphology of aging mitochondria in mutants defective or synthesis of ergosterol (*Δerg4*), phosphatidylserine (*Δcho1*), or cardiolipin (*Δcrd1*) appeared similar to wildtype cells (**FIG 3D-E**) indicating that these lipid species are dispensable for mitochondria morphology transitions in chronologically aging cells. Taken together, these experiments reveal that mitochondria morphology and sphingolipid metabolism are coupled in aging cells.

### Overexpression of Isc1 preserves mitochondria during chronological aging

SL catabolic activities in yeast include two ceramidases (Ydc1 and Ypc1) and an inositol phosphosphingolipid phospholipase (Isc1) homologous to sphingomyelinase enzymes in animal cells (Megyeri et al., 2016). We tested if loss of these catabolic activities affect mitochondria in chronologically aging yeast. Although mitochondrial morphology in strains deleted for any of these enzymes resembled wildtype cells, we observed a slight but significant increase in mitochondrial width in cells lacking Isc1 (**FIG 4A-B**). To test the hypothesis that accumulation of complex SLs contributes to altered mitochondrial morphology in aging cells, we generated yeast strains where the native promoter of each of these genes was swapped for the ADH1 promoter, which drives high levels of protein expression. While pADH1-driven expression of YPC1 and YDC1 had no affect on mitochondrial morphology in aging cells, pADH1-driven expression of ISC1 significantly reduced the width of mitochondria in chronologically aged cells (**FIG 4C-D**). These results suggest that accumulation of complex phospholipids (IPCs, MIPCs, and M(IP)_2_Cs) contributes to mitochondrial morphology transitions in aging cells, and increased Isc1 activity can prevent such changes.

**Figure 4.**
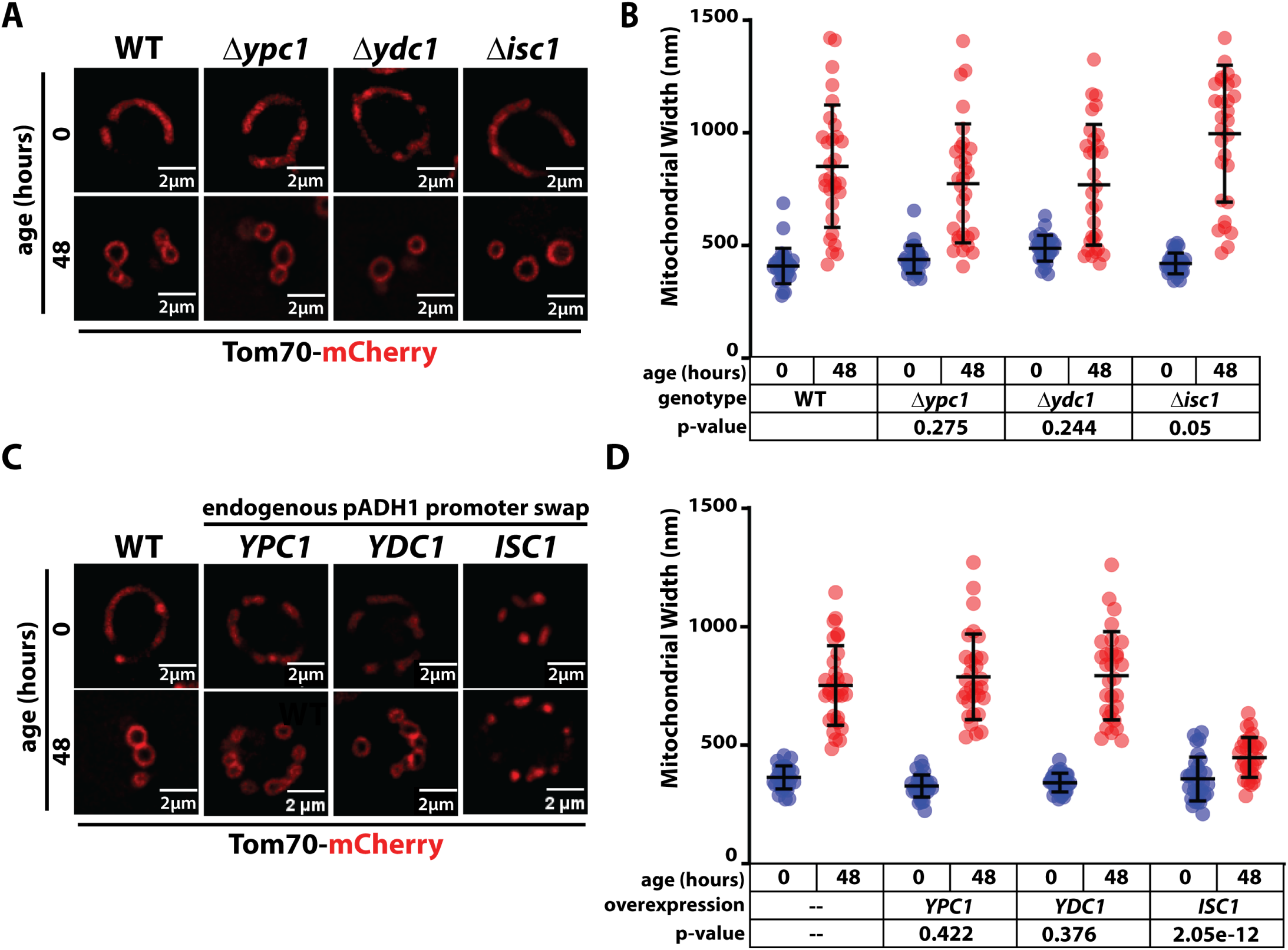
Isc1 modifies age-associated mitochondrial swelling. (A) Yeast expressing Tom70-mCherry were grown to mid-log phase (OD_600_ = 0.5) to initiate chronological aging time course (t=0). Fluorescence microscopy images were collected again after 48 hours of aging. Wildtype (WT) cells were analyzed as well as the indicated knockout mutants. (B) Mitochondria width was measured (n ≥ 30 cells; ±SD (error bars)) for each condition shown in (D). The p-value was computed using a paired T-TEST comparing the indicated mutant cells (t=48) to wildtype cells (t=48). (C) Yeast expressing Tom70-mCherry were grown to mid-log phase (OD_600_ = 0.5) to initiate chronological aging time course (t=0). Fluorescence microscopy images were collected again after 48 hours of aging. Wildtype (WT) cells were analyzed as well as the indicated overexpression strains. (D) Mitochondria width was measured (n ≥ 30 cells; ±SD (error bars)) for each condition shown in (D). The p-value was computed using a paired T-TEST comparing the indicated mutant cells (t=48) to wildtype cells (t=48).

### Oxidative stress alters mitochondrial morphology in an SL-dependent manner

Since yeast chronological aging is associated with oxidative stress (Chen et al., 2005; Longo et al., 2012) we tested if oxidative stress triggers mitochondrial morphology changes like those observed as cells age chronologically. We found that treatment of yeast with hydrogen peroxide (1, 2, and 3mM H_2_O_2_) triggered mitochondrial morphology changes similar to those observed in aging cells within 30 minutes of treatment (**FIG 5A-C** and **FIG S3A-C**). We also observed that pre-treatment of yeast cells with myriocin to deplete sphingolipids prior to hydrogen peroxide exposure significantly decreased these morphology changes to mitochondria (**FIG 5D-E**). Notably, the addition of PHS to the media abrogated the effects of myriocin treatment, indicating that exogenous uptake of phytoceramide is sufficient to restore the wider, rounded mitochondrial morphology associated with oxidative stress (**FIG 5D-E**). Finally, we found that pre-treatment of cells with myriocin to deplete sphingolipids promoted tolerance to oxidative stress (**FIG 5F** and **FIG S3D**). Taken together, these results reveal that SLs are required for mitochondrial morphology changes induced by acute oxidative stress. Furthermore, these SL-dependent changes in mitochondrial morphology correlate with sensitivity to oxidative stress.

**Figure 5.**
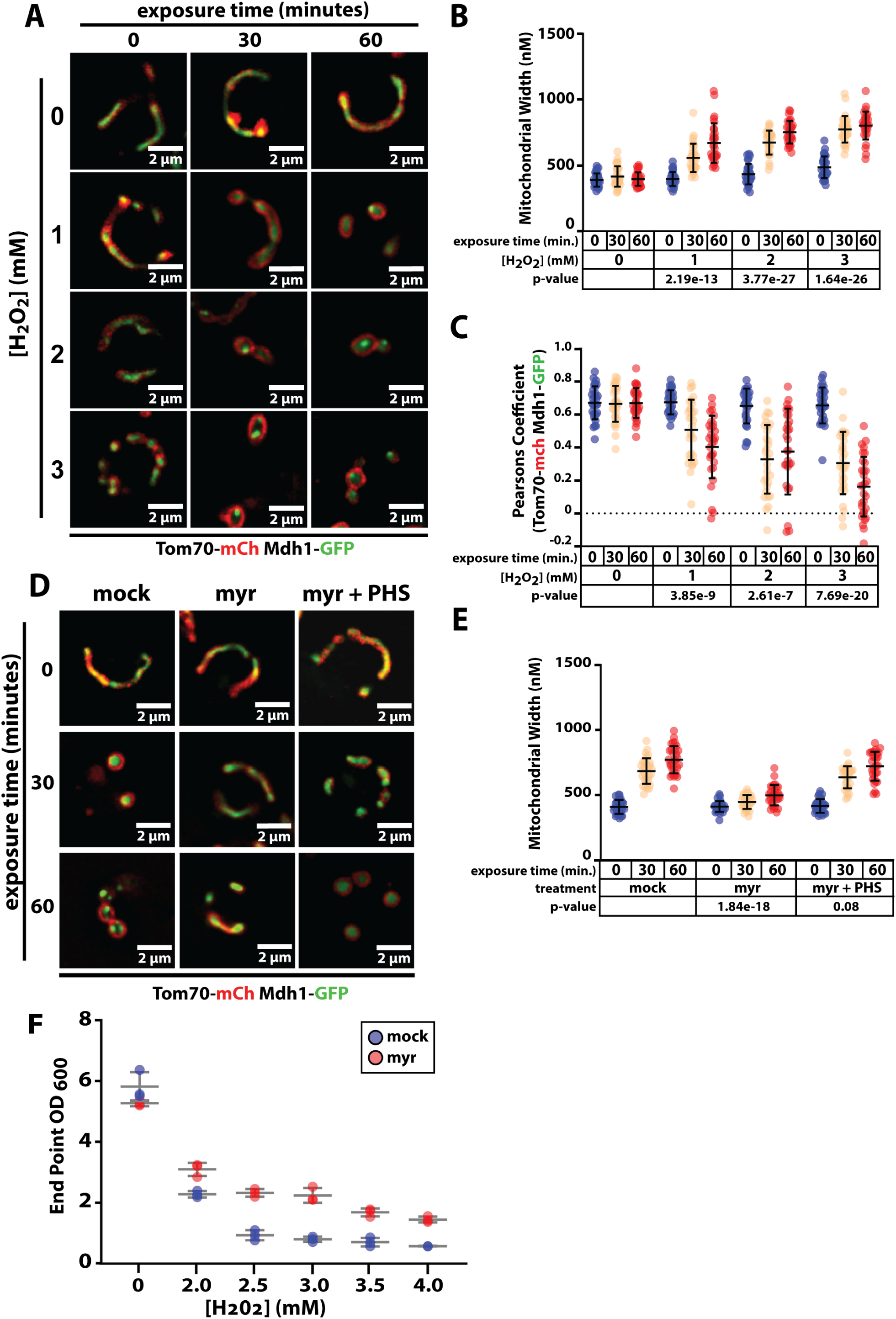
Sphingolipid-dependent mitochondria remodeling occurs during acute oxidative stress. (A) Yeast cells (strain background: SEY6210) expressing Tom70-mCherry (red) and Mdh1-GFP (green) were grown to mid-log phase (OD_600_ = 0.5) and subject to treatment with the indicated concentration of H_2_O_2_. Fluorescence microscopy images were collected at the indicated time points following exposure. (B) Mitochondria width was measured (n ≥ 30 cells; ±SD (error bars)) at each time point. The indicated p-value was computed using a paired T-TEST comparing the indicated time point to t=0. (C) Pearson correlation coefficient was computed for individual cells (n ≥ 30 cells; ±SD (error bars)) using Tom70-mCherry (red) and Mdh1-GFP (green) as markers of the outer mitochondrial membrane (OMM) and mitochondrial matrix (MM), respectively. The indicated p-value was computed using a paired T-TEST comparing the indicated time point to t=0. (D) Yeast expressing Tom70-mCherry (red) and Mdh1-GFP (green) were grown to mid-log phase (OD_600_ = 0.5) and subject to treatment with the indicated concentration of H_2_O_2_. Fluorescence microscopy images were collected at the indicated time points following exposure. (E) Mitochondria width was measured (n ≥ 30 cells; ±SD (error bars)) at each time point. The indicated p-value was computed using a paired T-TEST comparing the indicated time point to t=0. (F) Yeast cultures were grown to mid-log phase (OD_600_ = 0.5) and subject to treatment with the indicated concentration of H_2_O_2_ for 12 hours of growth before measuring OD_600_.

### Perspectives and future directions

Sphingolipids are structurally important lipids at the plasma membrane, although accumulating evidence indicates that they also have important functions in mitochondrial membranes (Jamil and Cowart, 2023; Spincemaille et al., 2014; Vos et al., 2021). Data presented in this study reveal a strong connection between sphingolipid metabolism and mitochondrial health in aging cells and during oxidative stress. We hypothesize that complex sphingolipid species may traffic to mitochondrial membranes during aging or oxidative stress, and the accumulation of SLs in mitochondrial membranes contributes to altered morphology and function. In conditions where sphingolipids are depleted, either pharmacologically or genetically, the decreased overall SL levels prevents remodeling of mitochondria normally observed during aging or oxidative stress. This also correlates with preserved MMP, suggesting that SL species may pose a specific liability for oxidative damage that can occur during environmental stress or aging. Indeed, accumulation of sphingolipids in mitochondria has been reported to occur in *pink1* mutant flies, a model of Parkinson’s Disease (Vos et al., 2021).

Although mitochondrial membranes are known to be enriched for specific SL species (Kitagaki et al., 2007), very little is known about how SL composition of mitochondria is managed and maintained. One clue may be that Isc1, the yeast homolog of sphingomyelinase enzymes, is reported to localize to mitochondria especially in stationary phase growth (Kitagaki et al., 2007; Vaena de Avalos et al., 2004). This suggests that Isc1 may be active on mitochondria membranes, where it could hydrolyze complex SL species (IPC, MIPC, and M(IP)_2_C) to produce localized pools of ceramides. Our finding that Isc1 overexpression prevents mitochondria remodeling in aging cells suggests that conversion of complex SLs to ceramides may protect and preserve mitochondria in aging cells. Consistent with these findings, *Δisc1* mutants exhibit both accumulation of ceramide species and phenotypes consistent with mitochondrial dysfunction (Kitagaki et al., 2007; Spincemaille et al., 2014). Together, these experiments suggest that Isc1 activity contributes to the maintenance of critical SL composition in mitochondrial membranes, which may become compromised during aging or stress.

These studies also raise important questions about the trafficking itinerary of complex sphingolipids, which are synthesized in the Golgi complex and accumulate in the plasma membrane of growing cells but may be become relocated to mitochondrial membranes during chronological aging or oxidative stress. One interesting avenue for future studies will involve dissecting the trafficking mechanisms that facilitate transport of complex SLs from peripheral membranes to mitochondrial membranes, which is likely to involve lipid transport proteins and inter-organelle contacts. The existence of such transport mechanisms would suggest that the observed changes in mitochondrial morphology represent a deliberate, adaptive remodeling response which may be part of a broader aging or stress response program. The fact that similar transitions are associated with aging and disease in other model organisms (García et al., 2013; Li et al., 2023; Rosa et al., 2023) suggests that this may be conserved across eukaryotic species, and that SL transport mechanisms discovered in yeast cells may be conserved in other eukaryotic organisms.

## Materials and methods

### Yeast strains and culture conditions

Yeast strains used in this study were derived from the SEY6210 background (*MATα leu2-3,112 ura3-52 his3-Δ200 trp1-Δ901 lys2-801 suc2-Δ9*) or the BY4741 (*MATa his3Δ1 leu2Δ0 met15Δ0 ura3Δ0*) background.

For chronological aging experiments, yeast strains were streaked from frozen stocks onto fresh Yeast Peptone Dextrose (YPD) solid media plates and incubated at 30°C. Single colonies were inoculated into 10mL YPD at 30°C with agitation at 220 rpm to an OD_600_ of 0.6. Yeast cells were then centrifuged at 4400 rpm for 1 minute and supernatant was discarded. Cells were washed with water and resuspended in SCD media and then subcultured into 10 ml of SCD at 0.3 OD. Cultures were returned to the incubator until OD_600_ of 0.6 was reached, which was considered to be the initial time point (t=0) for chronological aging experiments. Cultures were then incubated at 30°C (shaking at at 220 rpm) for the indicated number of hours.

### Fluorescence microscopy imaging

Fluorescence microscopy imaging of yeast cells was performed using a Delta Vision Elite Imaging system (Olympus IX-71 inverted microscope; Olympus 100× oil objective (1.4 NA); DV Elite sCMOS camera, GE Healthcare). Images obtained were deconvolved using SoftWoRx software (v7.0.0; GE Healthcare) and mean fluorescence intensity was measured using Fiji (NIH). For width measurements a line profile was used to measure the distance between the red (Alexa Fluor 594; 475 nm excitation) peaks of individual mitochondria. Colocalization analysis was performed using Pearson correlation coefficient measurement by the SoftWoRx software.

### Assessment of mitochondrial membrane potential

To assess mitochondrial membrane potential (MMP) we used the MitoLoc system ((Vowinckel et al., 2015)). In brief, the MitoLoc system uses a plasmid containing two fluorescently tagged proteins: preSu9-GFP (green), which localizes to mitochondria independent of MMP, and preCox4-mCherry (red) which localizes to mitochondria in a MMP-dependent manner. Thus, accumulation of mCherry signal outside of mitochondria indicates loss of MMP. Colocalization analysis of these two signals was performed using Pearson correlation coefficient measurement by the softWoRx software (v7.0.0; GE Healthcare).

### Analysis of Vacuolar Localization of Mitochondrial Markers

Vacuolar localization of mitochondrial proteins was assessed by fluorescence microscopy imaging of yeast cells expressing Tom70-mCherry and Mdh1-GFP as mitochondria markers and Vph1-mTagBPF2 as a marker of the vacuolar membrane. Collected images were deconvolved in SoftWoRx and analzyed in Fiji. Specifically, mean fluorescence intensity of Tom70-mCherry was measured within the vacuolar region of the cell, as marked by Vph1-mTagBFP2. Background fluorescence intensity was measured and subtracted to obtain the measurement of Tom70-mCherry localized to the vacuole.

### Analysis of Oxidative Stress Tolerance

To assess oxidative stress tolerance, cells were grown to mid-log phase in SCD media at 30°C in a shaking incubator (220 rpm). To initiate the assay cells were exposed to the indicated concentrations of hydrogen peroxide (H_2_O_2_) and allowed to continue to grow in the shaking incubator. After 12 hours of growth, OD_600_ was measured and recorded.

## Acknowledgements

We are grateful to T Graham, M Jimenez, and B Jain for technical advice and feedback. This project was supported by a Seeding Success Award from Vanderbilt University’s Office of Research and Innovation. ACE was supported by Vanderbilt’s Chemical Biology Interface Training Program (T32GM149371). TAM was supported by the Vanderbilt University VERTICES-PREP program (R25GM134979). KDF was supported by the Vanderbilt University MARC program (T34GM136451). NLH and JAM were supported by NIH grant R35 GM144112 (to JAM).

---

**Supplemental Figure S1.**
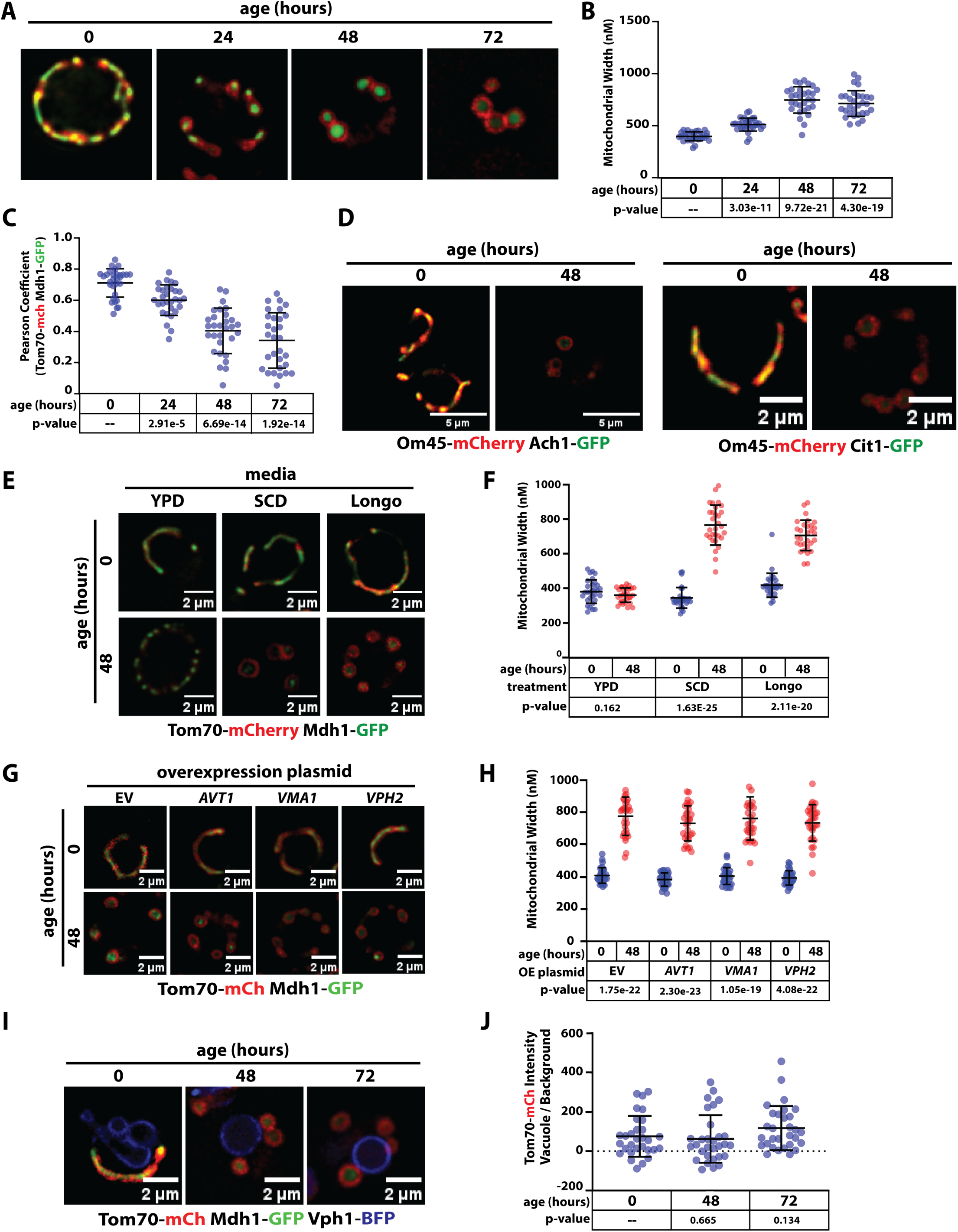
Characterization of mitochondria morphology changes in chronologically aging yeast cells. (A) Yeast cells (strain background: BY4741) expressing Tom70-mCherry (red) and Mdh1-GFP (green) were grown to mid-log phase (OD_600_ = 0.5) to initiate chronological aging time course (t=0). Fluorescence microscopy images were collected every 24 hours at the indicated time points. (B) Mitochondria width was measured (n ≥ 30 cells; ±SD (error bars)) for each time point shown in (A). The indicated p-value was computed using a paired T-TEST comparing the indicated time point to t=0. (C) Pearson correlation coefficient was computed for individual cells (n ≥ 30 cells; ±SD (error bars)) using Tom70-mCherry (red) and Mdh1-GFP (green) as markers of the outer mitochondrial membrane (OMM) and mitochondrial matrix (MM), respectively. The indicated p-value was computed using a paired T-TEST comparing the indicated time point to t=0. (D) Yeast cells (strain background: SEY6210) expressing various mitochondrial markers including Om45-mCherry (red), Ach1-GFP (green) and Cit1-GFP (green) were grown to mid-log phase (OD_600_ = 0.5) to initiate chronological aging time course (t=0). Fluorescence microscopy images were collected every 24 hours at the indicated time points. (E) Yeast cells (strain background: SEY6210) expressing Tom70-mCherry (red) and Mdh1- GFP (green) were grown to mid-log phase (OD_600_ = 0.5) to initiate chronological aging time course (t=0). Cells were then cultured in the indicated growth media and fluorescence microscopy images were collected every 24 hours at the indicated time points. (F) Mitochondria width was measured (n ≥ 30 cells; ±SD (error bars)) at each time point and condition shown in (F). The indicated p-value was computed using a paired T-TEST comparing the indicated time point to t=0. (G) Yeast cells (strain background: SEY6210) expressing Tom70-mCherry (red) and Mdh1- GFP (green) and harboring the indicated overexpression plasmid were grown to mid-log phase (OD_600_ = 0.5) to initiate chronological aging time course (t=0). Cells were then cultured in the indicated growth media and fluorescence microscopy images were collected every 24 hours at the indicated time points. (H) Mitochondria width was measured (n ≥ 30 cells; ±SD (error bars)) at each time point and condition shown in (H). The indicated p-value was computed using a paired T-TEST comparing the t=0 to t=48 for each indicated strain. (I) Yeast cells (strain background: SEY6210) expressing Tom70-mCherry (red) and Mdh1- GFP (green) and Vph1-mTagBFP2 (blue) were grown to mid-log phase (OD_600_ = 0.5) to initiate chronological aging time course (t=0). Fluorescence microscopy images were collected at the indicated time points. (J) Intensity of Tom70-mCherry signal in the vacuole lumen (normalized to background) was measured at each of the indicated time points. The indicated p-value was computed using a paired T-TEST comparing the t=0 to each indicated time point.

**Supplemental Figure S2.**
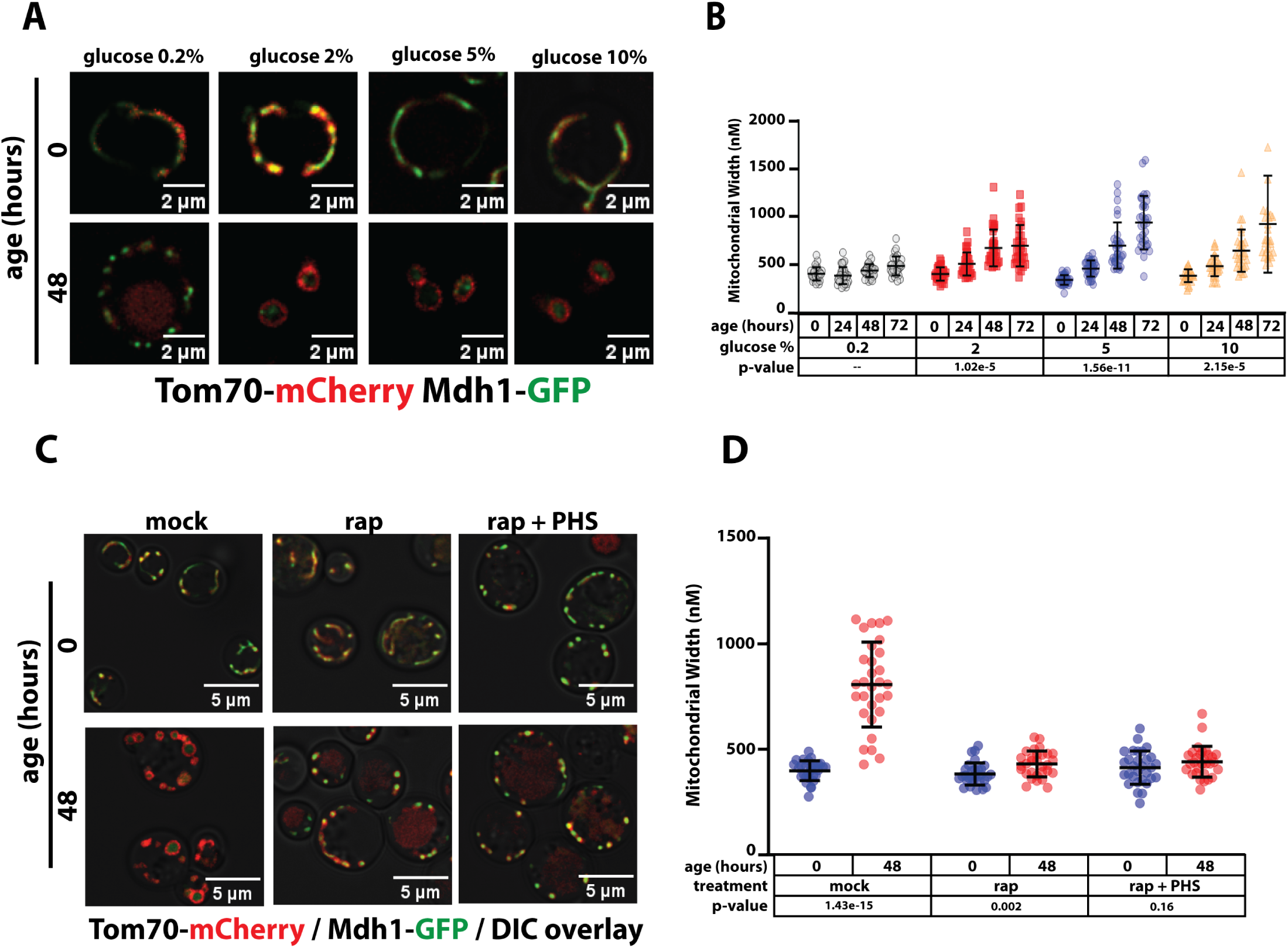
Interventions that modify mitochondrial morphology in chronologically aged yeast cells. (A) Yeast expressing Tom70-mCherry (red) and Mdh1-GFP (green) were grown to mid-log phase (OD_600_ = 0.5) to initiate chronological aging time course (t=0). Cells were then cultured in SCD media with the indicated glucose concentration and fluorescence microscopy images were collected at the indicated time points. (B) Mitochondria width was measured (n ≥ 30 cells; ±SD (error bars)) for each condition shown in (A). The indicated p-value was computed using a paired T-TEST comparing the indicated treatment (t=48) to mock treated cells (t=48). (C) Yeast expressing Tom70-mCherry (red) and Mdh1-GFP (green) were grown to mid-log phase (OD_600_ = 0.5) to initiate chronological aging time course (t=0). Cells were left untreated (mock) or treated with the following compounds associated with rapamycin (rap) or rapamycin + phytosphingosine (rap + PHS) and fluorescence microscopy images were collected at the indicated time points. (D) Mitochondria width was measured (n ≥ 30 cells; ±SD (error bars)) for each condition shown in (C). The indicated p-value was computed using a paired T-TEST comparing mid-log cells (t=0) to the aged cells (t=48).

**Supplemental Figure S3.**
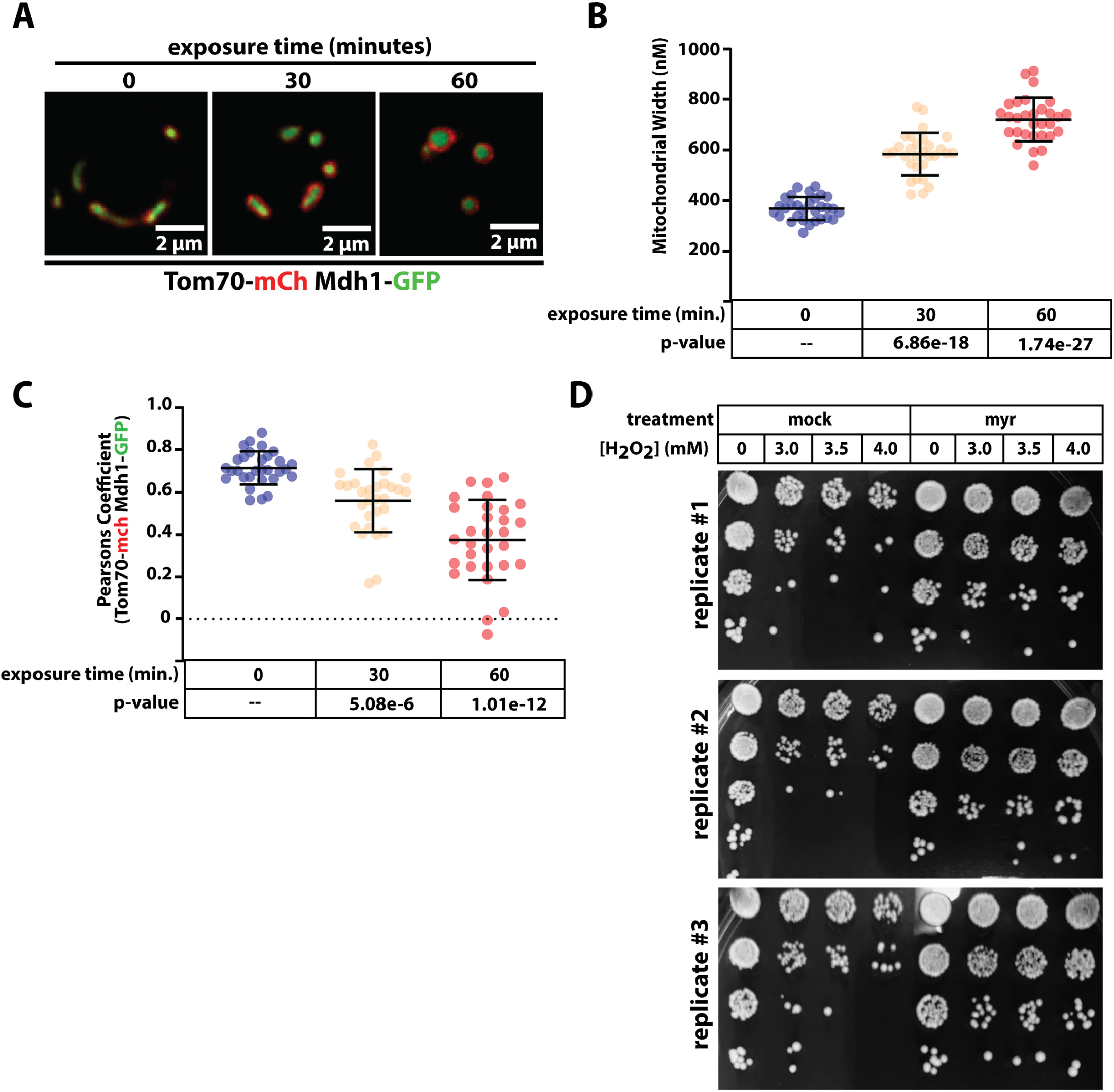
Sphingolipid-dependent mitochondria remodeling occurs during acute oxidative stress. (A) Yeast cells (strain background: BY4741) expressing Tom70-mCherry (red) and Mdh1-GFP (green) were grown to mid-log phase (OD_600_ = 0.5) and subject to treatment with the indicated concentration of H_2_O_2_. Fluorescence microscopy images were collected at the indicated time points following exposure. (B) Mitochondria width was measured (n ≥ 30 cells; ±SD (error bars)) at each time point. The indicated p-value was computed using a paired T-TEST comparing the indicated time point to t=0. (C) Pearson correlation coefficient was computed for individual cells (n ≥ 30 cells; ±SD (error bars)) using Tom70-mCherry (red) and Mdh1-GFP (green) as markers of the outer mitochondrial membrane (OMM) and mitochondrial matrix (MM), respectively. The indicated p-value was computed using a paired T-TEST comparing the indicated time point to t=0. (D) Yeast cultures were grown to mid-log phase (OD_600_ = 0.5) and subject to treatment with the indicated concentration of H_2_O_2_ for 12 hours of growth before and plating them on solid YPD media in two-fold serial dilutions.

## References

Chaudhari, S.N., and E.T. Kipreos. 2017. Increased mitochondrial fusion allows the survival of older animals in diverse C. elegans longevity pathways. Nat Commun. 8:182.

Chen, Q., Q. Ding, and J.N. Keller. 2005. The stationary phase model of aging in yeast for the study of oxidative stress and age-related neurodegeneration. Biogerontology. 6:1–13.

Cutler, R.G., K.W. Thompson, S. Camandola, K.T. Mack, and M.P. Mattson. 2014. Sphingolipid metabolism regulates development and lifespan in Caenorhabditis elegans. Mech Ageing Dev. 143–144:9-18.

García, M.L., A. Fernández, and M.T. Solas. 2013. Mitochondria, motor neurons and aging. J Neurol Sci. 330:18–26.

Gaudioso, A., P. Garcia-Rozas, M.J. Casarejos, O. Pastor, and J.A. Rodriguez-Navarro. 2019. Lipidomic Alterations in the Mitochondria of Aged Parkin Null Mice Relevant to Autophagy. Front Neurosci. 13:329.

Guo, J., X. Huang, L. Dou, M. Yan, T. Shen, W. Tang, and J. Li. 2022. Aging and aging-related diseases: from molecular mechanisms to interventions and treatments. Signal Transduct Target Ther. 7:391.

Hammerschmidt, P., D. Ostkotte, H. Nolte, M.J. Gerl, A. Jais, H.L. Brunner, H.G. Sprenger, M. Awazawa, H.T. Nicholls, S.M. Turpin-Nolan, T. Langer, M. Krüger, B. Brügger, and J.C. Brüning. 2019. CerS6-Derived Sphingolipids Interact with Mff and Promote Mitochondrial Fragmentation in Obesity. Cell. 177:1536–1552.e1523.

Hepowit, N.L., J.K.A. Macedo, L.E.A. Young, K. Liu, R.C. Sun, J.A. MacGurn, and R.C. Dickson. 2021. Enhancing lifespan of budding yeast by pharmacological lowering of amino acid pools. Aging (Albany NY). 13:7846–7871.

Houtkooper, R.H., L. Mouchiroud, D. Ryu, N. Moullan, E. Katsyuba, G. Knott, R.W. Williams, and J. Auwerx. 2013. Mitonuclear protein imbalance as a conserved longevity mechanism. Nature. 497:451–457.

Huang, X., B.R. Withers, and R.C. Dickson. 2014. Sphingolipids and lifespan regulation. Biochim Biophys Acta. 1841:657–664.

Hughes, A.L., and D.E. Gottschling. 2012. An early age increase in vacuolar pH limits mitochondrial function and lifespan in yeast. Nature. 492:261–265.

Hughes, A.L., C.E. Hughes, K.A. Henderson, N. Yazvenko, and D.E. Gottschling. 2016. Selective sorting and destruction of mitochondrial membrane proteins in aged yeast. Elife. 5.

Jamil, M., and L.A. Cowart. 2023. Sphingolipids in mitochondria-from function to disease. Front Cell Dev Biol. 11:1302472.

Kitagaki, H., L.A. Cowart, N. Matmati, S. Vaena de Avalos, S.A. Novgorodov, Y.H. Zeidan, J. Bielawski, L.M. Obeid, and Y.A. Hannun. 2007. Isc1 regulates sphingolipid metabolism in yeast mitochondria. Biochim Biophys Acta. 1768:2849–2861.

Li, A., G.J. Shami, L. Griffiths, S. Lal, H. Irving, and F. Braet. 2023. Giant mitochondria in cardiomyocytes: cellular architecture in health and disease. Basic Res Cardiol. 118:39.

Li, S., and H.E. Kim. 2021. Implications of Sphingolipids on Aging and Age-Related Diseases. Front Aging. 2:797320.

Lima, T.I., P.P. Laurila, M. Wohlwend, J.D. Morel, L.J.E. Goeminne, H. Li, M. Romani, X. Li, C.M. Oh, D. Park, S. Rodríguez-López, J. Ivanisevic, H. Gallart-Ayala, B. Crisol, F. Delort, S. Batonnet-Pichon, L.R. Silveira, L. Sankabattula Pavani Veera Venkata, A.K. Padala, S. Jain, and J. Auwerx. 2023. Inhibiting de novo ceramide synthesis restores mitochondrial and protein homeostasis in muscle aging. Sci Transl Med. 15:eade6509.

Longo, V.D., G.S. Shadel, M. Kaeberlein, and B. Kennedy. 2012. Replicative and chronological aging in Saccharomyces cerevisiae. Cell Metab. 16:18–31.

Megyeri, M., H. Riezman, M. Schuldiner, and A.H. Futerman. 2016. Making Sense of the Yeast Sphingolipid Pathway. J Mol Biol. 428:4765–4775.

Rosa, F.L.L., I.I.A. de Souza, G. Monnerat, A.C. Campos de Carvalho, and L. Maciel. 2023. Aging Triggers Mitochondrial Dysfunction in Mice. Int J Mol Sci. 24.

Sarnoski, E.A., P. Liu, and M. Acar. 2017. A High-Throughput Screen for Yeast Replicative Lifespan Identifies Lifespan-Extending Compounds. Cell Rep. 21:2639–2646.

Scheckhuber, C.Q., N. Erjavec, A. Tinazli, A. Hamann, T. Nyström, and H.D. Osiewacz. 2007. Reducing mitochondrial fission results in increased life span and fitness of two fungal ageing models. Nat Cell Biol. 9:99–105.

Singh, P., K. Gollapalli, S. Mangiola, D. Schranner, M.A. Yusuf, M. Chamoli, S.L. Shi, B. Lopes Bastos, T. Nair, A. Riermeier, E.M. Vayndorf, J.Z. Wu, A. Nilakhe, C.Q. Nguyen, M. Muir, M. G. Kiflezghi, A. Foulger, A. Junker, J. Devine, K. Sharan, S.J. Chinta, S. Rajput, A. Rane, P. Baumert, M. Schönfelder, F. Iavarone, G. di Lorenzo, S. Kumari, A. Gupta, R. Sarkar, C. Khyriem, A.S. Chawla, A. Sharma, N. Sarper, N. Chattopadhyay, B.K. Biswal, C. Settembre, P. Nagarajan, K.L. Targoff, M. Picard, S. Gupta, V. Velagapudi, A.T. Papenfuss, A. Kaya, M.G. Ferreira, B.K. Kennedy, J.K. Andersen, G.J. Lithgow, A.M. Ali, A. Mukhopadhyay, A. Palotie, G. Kastenmüller, M. Kaeberlein, H. Wackerhage, B. Pal, and V.K. Yadav. 2023. Taurine deficiency as a driver of aging. Science. 380:eabn9257.

Singh, P., and R. Li. 2018. Emerging roles for sphingolipids in cellular aging. Curr Genet. 64:761–767.

Spincemaille, P., N. Matmati, Y.A. Hannun, B.P. Cammue, and K. Thevissen. 2014. Sphingolipids and mitochondrial function in budding yeast. Biochim Biophys Acta. 1840:3131–3137.

Vaena de Avalos, S., Y. Okamoto, and Y.A. Hannun. 2004. Activation and localization of inositol phosphosphingolipid phospholipase C, Isc1p, to the mitochondria during growth of Saccharomyces cerevisiae. J Biol Chem. 279:11537-11545.

Vos, M., M. Dulovic-Mahlow, F. Mandik, L. Frese, Y. Kanana, S. Haissatou Diaw, J. Depperschmidt, C. Böhm, J. Rohr, T. Lohnau, I.R. König, and C. Klein. 2021. Ceramide accumulation induces mitophagy and impairs β-oxidation in PINK1 deficiency. Proc Natl Acad Sci U S A. 118.

Vowinckel, J., J. Hartl, R. Butler, and M. Ralser. 2015. MitoLoc: A method for the simultaneous quantification of mitochondrial network morphology and membrane potential in single cells. Mitochondrion. 24:77–86.

Zou, K., S. Rouskin, K. Dervishi, M.A. McCormick, A. Sasikumar, C. Deng, Z. Chen, M. Kaeberlein, R.B. Brem, M. Polymenis, B.K. Kennedy, J.S. Weissman, J. Zheng, Q. Ouyang, and H. Li. 2020. Life span extension by glucose restriction is abrogated by methionine supplementation: Cross-talk between glucose and methionine and implication of methionine as a key regulator of life span. Sci Adv. 6:eaba1306.

